# Genomic and Metabolic Hallmarks of SDH- and FH-Deficient Renal Cell Carcinomas

**DOI:** 10.1101/2021.06.09.445990

**Authors:** Angela Yoo, Cerise Tang, Mark Zucker, Kelly Fitzgerald, Phillip M Rappold, Kate Weiss, Benjamin Freeman, Chung-Han Lee, Nikolaus Schultz, Robert Motzer, Paul Russo, Jonathan Coleman, Victor E Reuter, Ying-Bei Chen, Maria I Carlo, Anthony J Gill, Ritesh R Kotecha, A. Ari Hakimi, Ed Reznik

## Abstract

**Purpose:** Succinate dehydrogenase-deficient and fumarate hydratase-deficient renal cell carcinomas (SDHRCC and FHRCC) are rare kidney cancers driven by loss of metabolically proximal enzymes. We sought to define and compare the genomic and metabolomic hallmarks of these entities.

**Experimental Design:** We analyzed SDHRCC and FHRCC tumors with either immunohistochemical evidence of loss of protein expression or genomically-confirmed biallelic inactivation of *SDHA/B/C/D/AF2* or *FH.* Somatic alterations were identified using clinical pipelines, and allele-specific copy number changes were identified using FACETS. Mass-spectrometry-based metabolomic profiling was performed on available SDHRCC and FHRCC tumors.

**Results:** Forty two patients were analyzed (25 FHRCC, 17 SDHRCC). In the germline analysis, 16/17 SDHRCC harbored a germline alteration in *SDHB,* whereas only 17/22 FHRCC had pathogenic germline *FH* variants. SDHRCC had a lower mutation burden (p = 0.02) and copy number alteration burden (p = 0.0002) than FHRCC. All SDHRCC presented with deletion of chromosome 1p (overlapping *SDHB*), whereas FHRCC demonstrated high but not ubiquitous loss of 1q (*FH* locus). Both SDHRCC and FHRCC demonstrated significant, idiopathic accumulation of the metabolite guanine. FHRCC tumors had elevated levels of urea cycle metabolites (argininosuccinate, citrulline, and fumarate), whereas SDHRCC had elevation of numerous acylcarnitines. These characteristic metabolic changes enabled the identification of a previously unrecognized SDH-deficient RCC.

**Conclusion:** Despite sharing similar genetic etiology, SDHRCC and FHRCC represent distinct molecular entities with unique genetic and metabolic abnormalities.

**Translational Relevance:** Mutations to the TCA cycle enzymes Succinate Dehydrogenase (SDH) and Fumarate Hydratase (FH) predispose individuals to unique subtypes of renal cell carcinoma (SDHRCC and FHRCC, respectively). Defining the genetic and metabolic hallmarks of these diseases is critical for advancing new diagnostic and therapeutic approaches for these rare but biologically intriguing entities. Despite a superficially similar genetic etiology, SDHRCC and FHRCC demonstrated significantly fewer secondary mutations to other cancer-associated genes and copy number aberrations than FHRCC, and was distinguished by universal loss-of-heterozygosity of chromosome 1p. Metabolomic analysis identified pathways disrupted in both SDHRCC and FHRCC, including the massive accumulation of free guanine, as well as pathways uniquely disrupted in each of the two entities. These metabolomic findings enabled the identification of a previously unidentified case of unclassified RCC with SDH deficiency, suggesting that metabolomic profiling may aid in phenotypic classification of tumors and uncover novel therapeutic targets.

## Introduction

Renal cell carcinomas (RCC) are a heterogeneous group of tumors which have become increasingly defined by morphologic and molecular features [1]. Despite significant diversity amongst RCC tumor subtypes, many share mutations in common genes integral to mitochondrial metabolism and oxidative phosphorylation *[2].* The 2016 WHO classification of renal tumors recognizes mutations in fumarate hydratase (encoded by the single gene *FH*) and succinate dehydrogenase (a protein complex composed of subunits encoded by *SDHA/SDHB/SDHC/SDHD* and assembled with *SDHAF2)* as distinct and rare histologic types of RCC (herein referred to as FHRCC and SDHRCC, respectively) [1]. Functionally, SDH and FH catalyze two consecutive and high-flux reactions in the TCA cycle, and SDH additionally encodes Complex II of the mitochondrial electron transport chain, which is responsible for oxidation of FADH_2_ [3]. Both SDHRCC and FHRCC are rare and it is speculated that SDHRCC constitutes 0.05%-0.2% of all RCC diagnoses [4], while FHRCC is comparatively more common but still rare, constituting 0.5-4.35% of all RCC diagnoses [5].

Due to their rarity, comprehensive evaluation of the genetic and metabolic features of SDHRCC and FHRCC tumors has been limited. *In vitro* data of mouse hepatocyte cell lines with silencing of *FH* or *SDHA* suggest that complete loss of either enzyme produces a block in the TCA cycle and consequent accumulation of related upstream metabolic intermediates *(i.e.* succinate in SDHRCC and fumarate in FHRCC) [6]. Both fumarate and succinate can act as potent inhibitors of alpha-ketoglutarate-dependent dioxygenases including DNA demethylases [7], and both have been implicated as drivers of hypermethylation phenotypes seen in RCC, paragangliomas, and pheochromocytomas [8], [9]. Both germline FHRCC and SDHRCC patients are predisposed to developing additional syndromic features; FHRCC patients can develop cutaneous and uterine leiomyomas [10], a constellation of genetic syndrome called hereditary leiomyomatosis and RCC (HLRCC); SDHRCC patients in contrast have increased risk of pheochromocytomas/paragangliomas [11] and gastrointestinal stromal tumors [12].

The functional similarity of the enzymes SDH and FH, and the similar downstream epigenetic changes caused by the accumulation of these oncometabolites, suggests that SDHRCC and FHRCC may bear genomic and metabolic similarities. Three recent studies, including one from our group, have described several genetic hallmarks of FHRCC, including characteristically low tumor mutation burden, distinct tumor microenvironments characterized by massive CD8+ T-cell infiltration with PD-L1+ expression in tumors, and a highly depleted and mutant mitochondrial genome [8,13,14]. Prior reports characterizing SDHRCC tumors have largely focused on clinicopathologic hallmarks of the disease and genetic analyses of specific *SDH* mutant alleles, but have not described the full genomic landscape of SDHRCC [5,15,16]. Hence, comprehensive molecular and metabolomic profiling of SDHRCC tumors has been limited [17].

To characterize and compare molecular characteristics of SDHRCC relative to FHRCC, we assembled a multi-institutional, genomically profiled cohort of SDHRCC and FHRCC tumors. For a subset of these tumors, we completed high-throughput semi-quantitative metabolomic profiling to identify characteristic and/or pathognomonic metabolic features of these entities. We hypothesized that because the substrates/products of SDH and FH are fundamentally connected to other physiologically critical metabolic pathways, SDHRCC and FHRCC may differ significantly in their metabolomic profiles. This focus on the metabolomic landscape of SDHRCC and FHRCC allowed us to identify metabolic phenotypes which separated TCA-cycle-mutant RCCs from clear cell RCC, and which further distinguished SDHRCC from FHRCC. We then applied metabolomic profiling as an exploratory exercise in a previously collected unclassified RCC dataset and identified a previously unrecognized SDHRCC patient within the cohort.

## Results

### Cohort Characteristics

We identified 83 RCC patients with a presumed diagnosis of FHRCC or SDHRCC from Memorial Sloan Kettering Cancer Center (MSK) and through a collaboration with a multi-institutional cohort [15], assembled from 15 institutions in North America, Europe, Asia, and Australia. The study was conducted under an IRB approved retrospective research protocol, including a waiver of consent. A combination of genomic, immunohistochemical (IHC), and expert genitourinary pathology review were used for case identification. The decision tree and criteria for the inclusion of each individual patient is provided in **Supplementary Figure 1** and **Supplementary Table 1**, respectively. Eleven patients were excluded due to lack of sequencing data (in the form of MSK-IMPACT [18], Whole Exome Sequencing (WES) or Whole Genome Sequencing (WGS)). Patients were subsequently included for analysis if they met either molecular criteria (biallelic loss of *FH* or *SDHA/SDHB/SDHC/SDHD/SDHAF2)* or IHC criteria (loss of FH expression and/or presence of 2-succino-cysteine (2SC) positive immunoreactivity [19], [20] or loss of SDHB expression [21]). When available, both germline and somatic data were integrated in order to determine biallelic status. Patients who did not meet these criteria (molecular or IHC) were excluded (n = 30). A total of 39 patients were included on the basis of genomic evidence and 3 patients were included under IHC criteria. The final cohort therefore consisted of a total of 42 patients: 25 patients were FH-deficient and 17 patients were SDH-deficient.

Patient and tumor characteristics are summarized in **Table 1.** Due to the multi-institutional nature of our cohort, a variety of sequencing approaches were completed on tumor specimens. All FHRCC patients were sequenced with MSK-IMPACT, an FDA-approved targeted sequencing assay which interrogates more than 341 cancer-associated genes including RCC-specific alterations such as *VHL, PBRM1, BAP1, SETD2, TP53, SDHA/B/C/D/AF2,* and *FH*. SDHRCC patients underwent multiple sequencing modalities including MSK-IMPACT, WES, and WGS. Three SDHRCC and two FHRCC patients had fresh frozen tumor tissue, some of which were untreated and some that received prior therapy, suitable for metabolomic analysis. These tumors were metabolomically profiled alongside frozen tumors from clear cell RCC (ccRCC, n= 14), unclassified RCC (URCC, n= 4), and adjacent normal kidney specimens (NK, n= 19).

SDHRCC and FHRCC patients presented with disease at a young age (median age 32, SDHRCC; median age 47 FHRCC, p = 0.03) and had comparable gender ratios. Other demographic and clinical information is provided in **Supplementary Table 1**. Of the 9 SDHRCC patients from our institution, 6 patients presented with localized disease (1 of whom went on to develop metastatic disease) and 3 SDHRCC patients were diagnosed with metastatic disease at initial presentation. There were 4 SDHRCC patients who received various systemic therapies; more information can be found in **Supplementary Table 2**. 12 FHRCC patients presented with localized disease whereas 13 patients presented with metastatic disease at initial presentation. Of the FHRCC patients who presented with localized disease, 11 patients experienced relapse. FHRCC patients received different types of systemic therapy, with combination therapy targeting both mTOR and VEGF as the most common and having the highest objective response rate [14].

### Distinct and recurrent copy number alterations are the primary events in the evolution of SDHRCC and FHRCC

To assemble a genomically defined cohort of SDHRCC with sufficient power to identify recurrent and discriminatory molecular features, we aggregated tumors profiled by different sequencing modalities into a multi-institutional cohort. Mutations and CNVs from all patients in the cohort are shown in **Supplementary Table 3** and **Supplementary Table 4**, respectively. The SDHRCC portion of our cohort was composed of 11 patients with matched tumor and normal *(e.g.* blood) sequencing data, of which 7 were profiled with MSK-IMPACT, 2 were profiled with WES, and 2 were profiled with WGS. Six additional patients from an outside institution had WES of tumor only (normal tissue was of insufficient quality for sequencing) and were used for validation of specific findings where possible, specifically the occurrence of loss of heterozygosity (LOH) on the p arm of chromosome 1 in SDHRCC and the presence of biallelic alterations. All FHRCC patients were profiled by MSK-IMPACT.

We first focused on identifying recurrent mutations and copy number alterations which define SDHRCC, and contrasting them with the corresponding genomic features in FHRCC. We identified biallelic alterations in all 17 patients with SDHRCC, whereas in FHRCC biallelic alterations were evident in 22 out of 25 patients. 16 out of 17 SDHRCC patients had pathogenic germline mutations; one SDHRCC patient sequenced with WGS had biallelic *SDHB* inactivation via somatic mutation and concomitant LOH (**Figure 1a**). Interestingly, the somatic mutation observed in this patient (at amino acid 242) has been previously reported as a germline mutation, suggesting evolutionary convergence on deleterious alleles [22]. FHRCC patients demonstrated a lower rate of germline alterations: 17 out of 22 FH patients who had consented to germline analysis had pathogenic germline mutations. This suggests that the majority of SDHRCC and FHRCC cases are associated with germline alterations, but both entities can arise through purely somatic biallelic inactivation.

**Figure 1.**
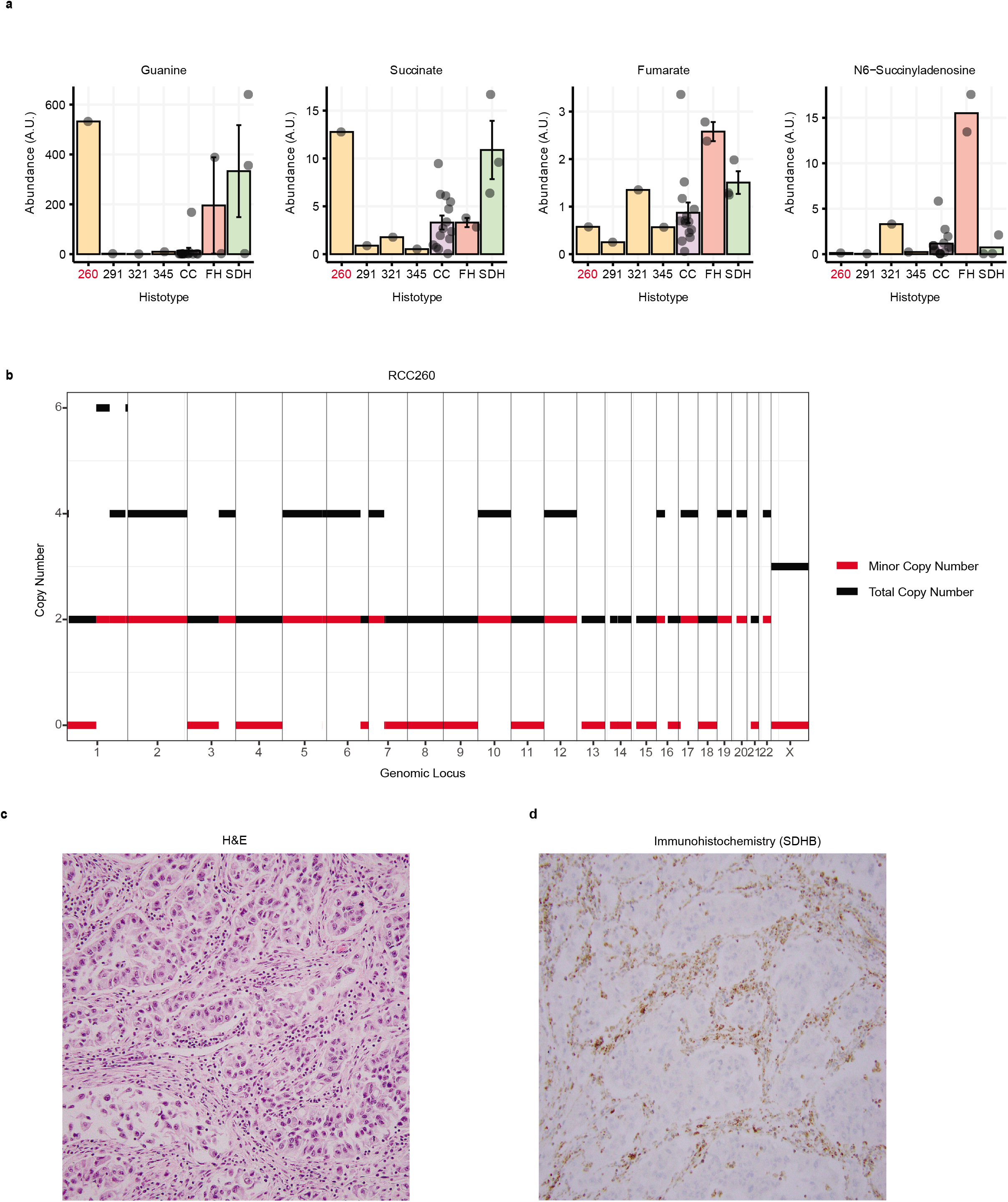
Genomic analysis and comparison of FHRCC and SDHRCC. (A) Comparison of incidence of germline mutations present in the FHRCC cohort vs. SDHRCC cohort. SDHRCC tumors are more likely to harbor pathogenic germline variants. Only 22 out of 25 FHRCC patients were consented for germline mutation analysis. (B) FHRCC tumors have a higher tumor mutation burden and (C) fraction of the genome altered (D) Oncoprint displaying recurrent (genes mutated in ≥3 patients) somatic mutations in FHRCC and SDHRCC tumors. Within the oncoprint, 2 out of 11 SDHRCC and 10 out of 25 FHRCC had recurrent mutations. However, 5/11 SDHRCC (45%) and 21/25 (84%) FHRCC when evaluating any occurrence of at least one somatic mutation. (E) Copy number profiles of FHRCC and SDHRCC. Top panels indicate the fraction of samples with gains. Bottom panel indicates the fraction of samples with LOH (including copy-neutral LOH). SDHRCC demonstrates universal LOH of chromosome arm 1p, whereas FHRCC often demonstrates LOH of 1q.

We next evaluated whether secondary somatic mutations in other cancer-associated genes present in SDHRCC and FHRCC converged on common molecular pathways. We first calculated coarse-grained measures of the copy number alteration burden (FGA, fraction genome altered, *i.e.* the fraction of the genome which was not diploid and heterozygous) and total somatic mutation burden (TMB) between FHRCC and SDHRCC patients. We restricted calculation of TMB to MSK-IMPACT 341 genes because they constituted the overlapping set of genes sequenced across all samples in our cohort. FHRCC tumors were in general more highly mutated than SDHRCC tumors, with significantly higher TMB (p = 0.02, **Figure 1b**), and had significantly more chromosomal copy number aberrations (FGA, p = 0.0002, Wilcoxon test, **Figure 1c**). In our efforts to genomically characterize FHRCC, we observed that the most commonly mutated genes in FHRCC patients, after *FH,* were *NF2* (n=5)*, FAT1* (n=3)*, PTPRT* (n=3), and *EP300* (n=3) **(Figure 1d)**. Notably, these comparatively commonly mutated genes are all tumor suppressor genes. Four of the five *NF2* mutations present are approximately clonal (cancer cell fraction (CCF) > 0.75), while two out of three *FAT1, PTPRT,* and *EP300* mutations each appear to be approximately clonal. Most of these mutations (all *NF2* mutations, 2 of 3 *FAT1* mutations, 2 of 3 *EP300* mutations, and 1 of 3 *PTPRT* mutations) were accompanied by LOH and therefore represent *bona fide* biallelic loss of these classical tumor suppressors. SDHRCC tended to have few, if any, accompanying somatic mutations at all, with a single case demonstrating an *NF2* mutation, which appeared to be subclonal. It was particularly notable that both SDHRCC and FHRCC lacked a significant number of mutations in common genes involved with ccRCC including *PBRM1* (n=1)*, SETD2* (n=0), *TERT* (n=1), and *TP53* (n=1). In sum, while the overall mutation burden across SDHRCC and FHRCC tumors was low, FHRCC tumors comparatively had higher TMB, FGA, and more often exhibited biallelic co-mutation of several additional tumor suppressor genes compared to SDHRCC.

In contrast to somatic mutations, SDHRCC and FHRCC demonstrated numerous large-scale copy number alterations (CNAs, **Figure 1e**). Although SDHRCC patients generally possessed fewer CNAs, all exhibited LOH (either copy number loss or copy-neutral LOH) on the p-arm of chromosome 1. This region overlaps with the *SDHB* locus, though typically the whole arm was lost. Notably, all 6 SDHRCC patients who received WES without matched normal (*e.g.* blood) tissues demonstrated LOH on 1p on manual inspection. In ~50% of cases, LOH of 1p was accompanied by concomitant single-copy amplification of 1q. Outside of chromosome 1, there were no recurrent CNVs among the SDHRCC patients. FHRCC patients in contrast usually – though not always – had LOH on 1q, particularly in the region overlapping *FH*. FHRCC showed higher incidence of other CNVs, including gains on chromosomes 16 and 17, which occur in >40% of FHRCC patients. FHRCC patients were also more likely to have gains on chromosome 2, with about one third having this alteration, while only 1/11 SDHRCC patients had a gain on chromosome 3.

### Distinct metabolic phenotypes beyond the TCA cycle underlie SDHRCC and FHRCC

Complete loss of SDH and FH *in vitro* causes the accumulation of the upstream metabolites succinate and fumarate, respectively. Other critical metabolic pathways, including the mitochondrial electron transport chain and the urea cycle, draw on or contribute to fumarate and succinate pools, rendering these pathways sensitive to large fluctuations in the concentrations of TCA cycle metabolites. To study the metabolic alterations underlying SDHRCC and FHRCC *in vivo,* we identified tumors from our genomics cohort with suitable fresh-frozen tissue for metabolomic profiling. Due to both the rarity of these diseases and the requirements for frozen tissue, our metabolomics cohort consisted of 3 SDHRCC and 2 FHRCC tumors. To contextualize the metabolomic profiles of these tumors, we also included 14 ccRCC samples, 4 unclassified RCC (URCC) samples, and 19 matched normal kidney (NK) tissue samples. In total, we quantified 600 metabolites across these samples. Principal components analysis (PCA) separated FH/SDH-deficient RCC both from matched normal tissue and from ccRCC, indicating that these entities broadly display unique metabolic phenotypes relative to ccRCC (**Figure 2a**). Importantly, we observed no separation between normal samples derived from SDH/FHRCC and ccRCC patients, respectively, indicating that these two diseases do not induce distinct metabolic changes in nearby kidney tissue and that *FH* and *SDHB* are metabolically haplosufficient in the kidney.

**Figure 2.**
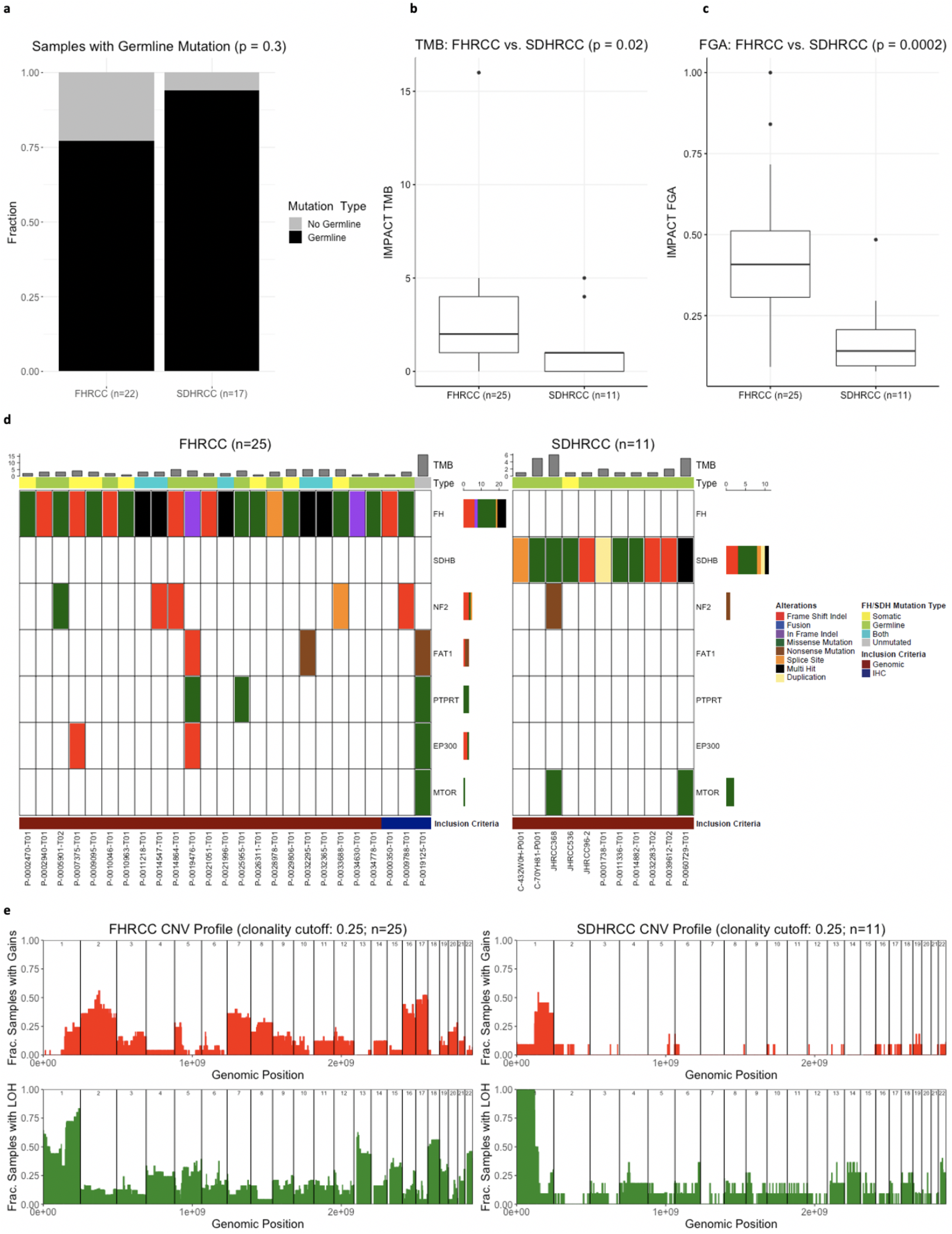
Metabolomic analysis and comparison of FHRCC and SDHRCC. (A): PCA plot of ccRCC (n=14), Unclassified tumors (n=4), SDHRCC (n=3) FHRCC tumors (n=2). The unclassified RCC tumors as well as the SDHRCC and FHRCC tumors cluster away from the clear cell tumors. (B) Barplot showing levels of succinate, fumarate, and guanine in normal kidney tissue, clear cell RCC, FHRCC, and SDHRCC tumors. (C) Volcano plot of metabolites that were elevated in SDHRCC/ FHRCC tumors compared to normal tissue, including succinate and guanine. (D) Urea cycle metabolites (argininosuccinate and citrulline) are more elevated in FHRCC vs normal as compared to SDHRCC vs normal. AGS (argininosuccinate), Fum (fumarate), Arg (arginine), Ure (urea), Orn (ornithine), Cit (Citrulline), Asp (aspartate) (E) Barplot showing levels of argininosuccinate, urea, citrulline, and arginine in normal kidney tissue, clear cell, FHRCC, and SDHRCC tumors. Argininosuccinate and citrulline were uniquely elevated in FHRCC but urea and arginine were not. (F) Barplot depiction of various acylcarnitine expression levels in SDHRCC compared to FHRCC that demonstrate an elevation of acylcarnitine in SDHRCC tumors.

### TCA cycle intermediates differ in abundance between FH- and SDH-deficient cancers

We noted that of the 8 quantified TCA cycle metabolites, succinate demonstrated the largest increase in abundance compared to normal tissue when grouping FHRCC and SDHRCC samples together, but succinate accumulation was more substantial in SDHRCC (log_2_fold-change 3.41, p-value = 0.001, q-value = 0.08, **Supplementary Table 5, Figure 2b**) than in FHRCC (log_2_fold-change 1.69, p-value = 0.02, q-value = 0.34). In contrast, fumarate levels were uniquely elevated in FHRCC, consistent with the position of SDH upstream of FH. Despite complete loss of *FH,* FHRCCs demonstrated only a ~2fold increase (p-value = 0.02, q-value = 0.34) in fumarate levels, in contrast to the ~50 fold accumulation observed when *FH* is ablated *in vitro [23].* Due to the limitations of metabolomic profiling of bulk tumor samples, we were unable to determine if this effect is due to the presence of infiltrating stromal/immune cells with normal fumarate levels, increased mechanisms for fumarate detoxification *in vivo,* or if FHRCC tumors themselves are at characteristically lower levels of fumarate than expected from cell line data.

### Guanine metabolism is impacted, but other purine metabolism is preserved, in both SDHRCC and FHRCC

We next examined in detail the metabolic consequences of SDH or FH loss beyond the TCA cycle. We identified 152 metabolites (excluding succinate) demonstrating statistically significant differential abundance between FH/SDH-RCC and normal kidney (q <0.2 and log_2_ fold change > 0.5, **Figure 2c**). Among the metabolomic adaptations in SDHRCC and FHRCC, the purine derivative guanine was observed to have the largest fold-change (~250-fold; 7.97 log_2_ fold change, p-value 0.002, q-value 0.04, **Figure 2b**). Interestingly, extreme elevation of guanine in FHRCC and SDHRCC was accompanied by a smaller elevation in guanosine (1.57 log_2_ fold change, p-value 0.0002, q-value 0.01), but no change in purine-derived adenine (−1.24 log_2_ fold change, p-value 0.16, qvalue 0.33) or adenosine (−1.42 log_2_ fold change, p-value 0.12, qvalue 0.30), and no change in the catabolic product of guanine, xanthine (−0.82 log_2_ fold change, p-value 0.088, qvalue 0.24). Importantly, guanine was also elevated ~20-fold relative to ccRCC tumors (p-value 0.005, q-value 0.11, **Figure 2b**), indicating that this metabolic phenotype was specific to disruption of FH and SDH. Levels of succinate, fumarate and guanine in normal kidney tissue, ccRCC, SDHRCC and FHRCC are compared in **Figure 2b**. To further confirm that the elevation of guanine was indicative of disruption of TCA cycle enzymes, we identified and metabolomically profiled 6 tumor regions and 3 normal regions from an additional FHRCC tumor. We again observed elevation of guanine (8.60 log_2_ fold change relative to normal, p-value 0.02, q-value 0.05, **Supplementary Table 6**) and guanosine (1.14 log_2_ fold change relative to normal, p-value 0.02, q-value 0.05), no elevation of adenine (0.66 log_2_ fold change relative to normal, p-value 0.17, q-value 0.24) or xanthine (−0.43 log_2_ fold change relative to normal, p-value 0.55, q-value 0.62), and a small elevation of adenosine (1.32 log_2_ fold change, p-value 0.02, q-value 0.05). The extremely large elevation of the guanine pool in SDHRCC and FHRCC suggests that loss of either FH or SDH may cause either overflow into free guanine from a peripheral pathway, or alternatively may prevent the turnover of free guanine by enzymes that rely on an intact TCA cycle.

### Urea cycle metabolites and acylcarnitine metabolism are differentially altered in FH- and SDH-deficient cancers

Although FHRCC and SDHRCC displayed common metabolic alterations, we noted that the two entities nevertheless clustered separately by PCA, indicating that they were characterized by qualitatively different metabolic phenotypes. Motivated by the statistical limitations of our small sample size, we examined our data for modules of physiologically-related metabolites demonstrating large differences in abundance between FHRCC and SDHRCC. Consistent with prior reports [23,24], we found that several metabolites in the urea cycle were specifically elevated in FHRCC but not SDHRCC (**Figure 2d**) when compared to normal kidney tissue, including argininosuccinate (log_2_ fold-change 1.21, p-value = 0.06, q-value = 0.34 in FHRCC; log_2_ fold-change −0.96, p-value = 0.34, q-value = 0.56 in SDHRCC) and citrulline (log_2_ fold-change 1.94, p-value = 0.03, q-value = 0.34 in FHRCC; log2 fold-change 0.80, p-value = 0.10, q-value = 0.37 in SDHRCC). Other urea cycle metabolites including arginine (log_2_ fold-change 0.15, p-value = 0.77, q-value = 0.90 in FHRCC; log_2_ fold-change 0.02, p-value = 0.46, q-value = 0.64 in SDHRCC) and urea (log_2_ fold-change −0.65, p-value = 0.40, q-value = 0.66 in FHRCC; log_2_ fold-change 0.03, p-value = 0.72, q-value = 0.82 in SDHRCC) did not show significant changes in abundance (**Figure 2e**). This is likely because they are downstream of the enzymatic reversal argininosuccinate lyase that occurs in the presence of excess fumarate. Increased urea cycle metabolites in FHRCC are consistent with previous reports of fumarate shunting to the urea cycle [23,24]. Excess fumarate is detoxified in the urea cycle by forming argininosuccinate, which can then be excreted into the urine.

In addition to the urea cycle, we identified several other functionally related groups of metabolites which distinguished FHRCC from SDHRCC. Several of the metabolites which were uniquely elevated in FHRCC when compared to SDHRCC appeared to be driven by inhibition of fumarate-producing reactions. The metabolite with the largest effect size was N6-succinyladenosine (~4.38 log_2_ fold, p-value 0.2). This metabolite commonly accumulates in the plasma of patients with germline fumarase deficiency, presumably as a result of excess of fumarate, which makes the production of fumarate and adenosine monophosphate (AMP) from N6-succinyladenosine unfavorable. Conversely, we also searched for metabolites whose accumulation was unique to SDHRCC, but not FHRCC. Intriguingly, we found that accumulation of acylcarnitines arose specifically in SDHRCC but not FHRCC, ccRCC, or adjacent normal tissues. Of the 33 acylcarnitine species measured in our dataset, we found 28/33 were more elevated in SDHRCC compared to FHRCC (**Figure 2f**). Mechanistically, acylcarnitines are intermediates of the FAD-dependent beta oxidation of fatty acids. Because Complex II of the electron transport chain is responsible for a significant amount of FAD regeneration, the accumulation of acylcarnitines specifically in SDHRCC suggests that Complex II deficiency impairs normal fatty acid oxidation.

### Differential metabolic signatures can be used to distinguish SDH vs FH deficiency in a diagnostically inconclusive patient

As noted earlier, PCA analysis indicated that the 4 URCC tumors included in our metabolomics data clustered closely with 3 SDHRCC and 2 FHRCC tumors, but separately from ccRCC (**Figure 2a**). Although this suggests that some URCC may metabolically resemble non-clear cell histologies such as SDHRCC and FHRCC, we also considered the possibility that some of the included URCC tumors in this small cohort may in fact harbor a previously overlooked loss of FH or SDH. Reasoning that idiopathic accumulation of guanine, in combination with succinate, fumarate, and n6-succinyladenosine, strongly distinguished SDHRCC/FHRCC from ccRCC and normal kidney, we investigated the levels of these metabolites in the 4 URCC samples. Only one tumor (RCC260) demonstrated elevated guanine (log_2_fold-change 5.30) and succinate (log_2_fold-change 1.95), but not fumarate (log_2_fold-change −0.60) or n6-succinyladenosine (log_2_fold-change −3.18) when compared to ccRCC (**Figure 3a)**. This suggested that RCC260 may be SDH-deficient. For 3/4 URCC tumors (excluding RCC345), we had matched MSK-IMPACT sequencing. Of these, only RCC260 demonstrated arm-level loss of chromosome 1p (**Figure 3b**), which was a universal event in SDHRCC.

**Figure 3.**
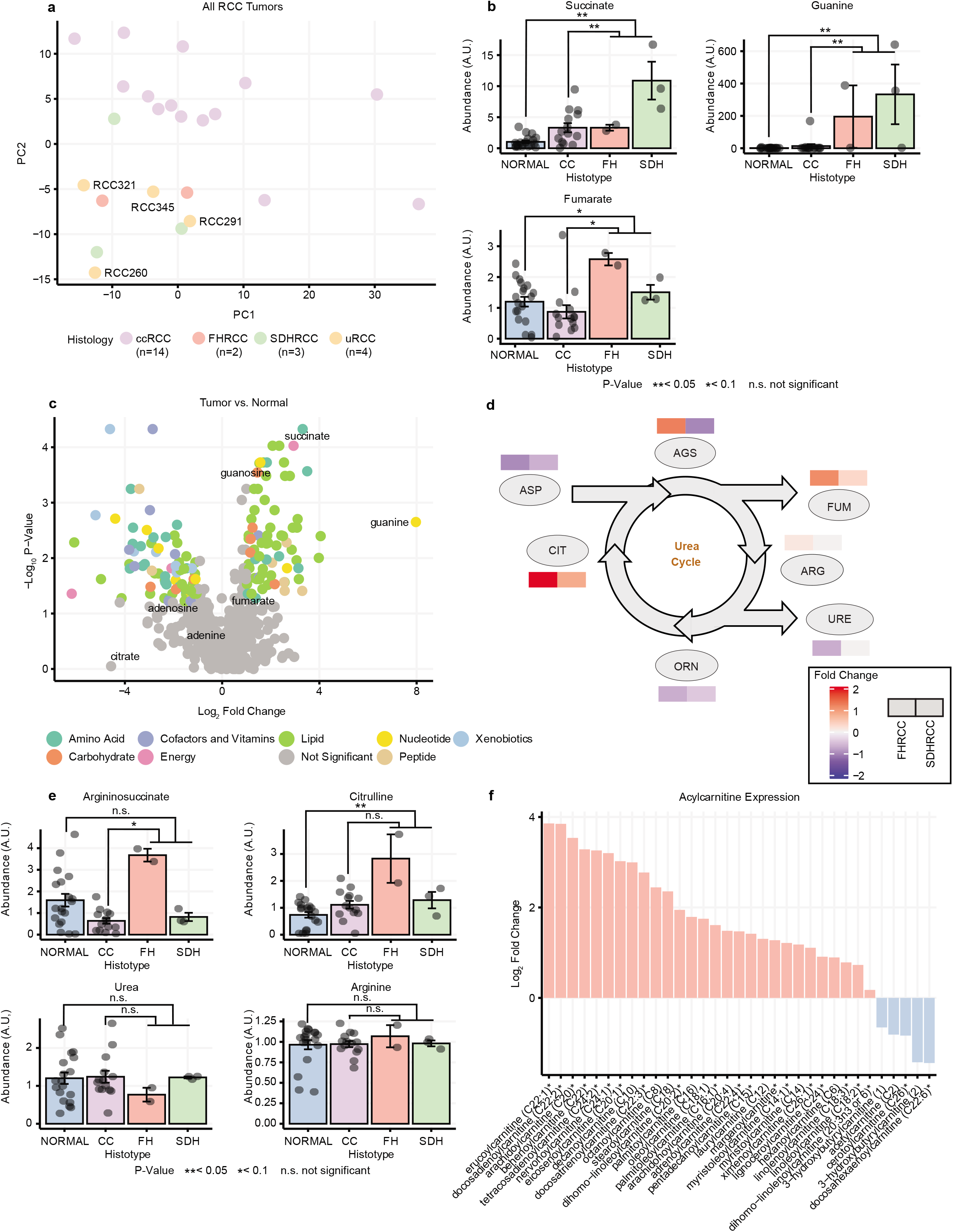
Metabolomic comparison of Unknown Sample with FHRCC and SDHRCC. Barplot of levels of guanine, succinate, fumarate, and N6-Succinyladenosine in four unclassified RCC samples [260, 291, 321, 345], ccRCC, FHRCC, and SDHRCC tumors. Sample 260 demonstrated extreme elevation of guanine and succinate, without elevation of fumarate, and N6-succinyladenosine. This resembles the metabolic profile of other SDHRCC tumors. (B) Copy Number profile of sample RCC260. Black indicates total copy number and red indicates the minor copy number. Note copy-neutral LOH of chromosome arm 1p, the locus of *SDHB* gene (C) Representative H&E image of the tumor RCC260 showing infiltrating tubules and nests of neoplastic cells with high grade nuclear features, eosinophilic cytoplasm, and scattered cytoplasmic vacuoles. (D) SDHB immunohistochemical stain of RCC260 shows a loss of SDHB protein expression in the neoplastic cells, whereas the stain is retained in the stromal, endothelial, and inflammatory cells.

Patient RCC260 was a 44 year old male patient who presented with anemia and was discovered to have a large kidney mass measuring 14.6 cm on imaging along with multiple bone lesions. Dedicated genitourinary pathology review of a bone biopsy noted a high grade, unclassified RCC subtype with suspicion of distal nephron differentiation. He later was treated with first-line pazopanib and ultimately passed 10 months later. Critically, this patient was among the first to be consented to MSK-IMPACT sequencing through our institution’s prospective clinical sequencing protocol, several years before SDHRCC and FHRCC were recognized as distinct entities. There was no evidence of somatic mutation of either *SDH* or *FH* via MSK-IMPACT testing. Germline status of FH and SDH subunits was unknown. Given the presence of 1p LOH and the characteristic metabolic features seen, we hypothesized that RCC260 may be a SDHRCC tumor. To test this hypothesis, we performed H&E and IHC staining of *SDHB* in RCC260. IHC analysis showed loss of *SDHB* protein expression in the neoplastic cells, but retention in stromal, endothelial, and inflammatory cells (**Figures 3c, 3d)**. Genomic, metabolomic, and IHC evidence therefore suggests that RCC260 represents a SDHRCC case likely with an unidentified germline alteration. Together, the data above argues that metabolomic profiling and germline testing (especially in URCC tumors) may be valuable supportive evidence in the diagnosis of SDHRCC and FHRCC.

## Discussion

We present here a comprehensive genomic and metabolomic analysis of SDH- and FH-deficient RCC tumors. To the best of our knowledge, this multi-institutional cohort is the largest compilation of SDHRCC patients with in-depth molecular correlates. The central finding of this analysis is that, despite superficial similarities in their genetic etiology, SDHRCC and FHRCC are genomically and metabolically distinct entities. SDHRCC tumors are comparatively genomically quiet, universally harbor 1p loss-of-heterozygosity, and rarely accrue co-mutations in cancer-associated genes. Both SDHRCC and FHRCC idiopathically accumulate free guanine, in addition to disease-specific alterations in acylcarnitine and urea cycle metabolites, respectively. Leveraging these molecular features, we retrospectively identified and immunohistochemically confirmed SDH deficiency in a previously unclassified RCC patient, highlighting the utility of metabolomics as an additional tool to distinguish ambiguous RCC variants.

Comparing SDHRCC and FHRCC patients, we found differences in clinical presentations. In our cohort, the median age at diagnosis of SDHRCC was 32 (range 15-68 years) which remains in line with prior reports (noting that 3 patients in a prior study were also included in this SDHRCC cohort) [4]. The male to female ratio was similar in both diseases, with males being the predominant gender (71% for SDHRCC and 65% for FHRCC). Nine of 27 (33%) patients in the prior study of SDHRCC [4] developed metastatic disease, 2 after prolonged follow-up of 5.5 and 30 years. In our MSKCC-restricted cohort with median follow-up of 21 months, 3 out of 9 SDHRCC patients presented with metastatic disease; of the remaining 6 patients with follow-up data, 1 patient developed metastatic disease. For FHRCC patients, our cohort was similar to previously described FHRCC cohorts [25]. FHRCC patients at our institution were diagnosed at a median age of 47 years, with a range of 20-74 years. Fourteen FHRCC patients were alive and twelve were deceased, again in concordance with the proportion of patients who were alive and deceased in a previous report of 32 FHRCC patients [25].

Both SDHRCC and FHRCC may be associated with syndromic features, and we assessed personal and family history for associated disease manifestations. Considering only patients from our own institution for which we could examine the full medical record in detail, 3 out of 9 SDHRCC patients had personal history of syndromic features (2 had pheochromocytomas and 1 had a paraganglioma); 3 out of 9 SDHRCC patients had a family history of kidney cancer. FHRCC patients from our institution also displayed syndromic features in their personal history such as cutaneous leiomyomas (n=1), uterine fibroids (n=4), and a combination of uterine fibroids and cutaneous leiomyomas (n = 2). More than half the FHRCC patients (n=14) did not demonstrate a family history of syndromic features. Summarized information can be found in **Table 1** while further details pertaining to each patient can be found in **Supplemental Table 1.** These data support the notion that “FH-deficient RCC” is preferred over “HLRCC-associated RCC” in order to encompass tumors with compatible histology and IHC evidence of FH-deficiency (FH loss and/or 2SC reactivity), but with uncertain clinical and family history. As the majority of our cohort did not have associated manifestations seen within a syndromic background despite enrichment for germline loss, this emphasizes that FHRCC and SDHRCC can be suspected in patients without family history as well. Of note, we also did not observe any *SDHA* germline mutations in our SDHRCC cohort. We observed that nearly all (16/17) SDHRCC patients exhibited germline mutations, consistent with a prior study that found germline variants in all 17 patients who underwent genetic testing (16 in *SDHB* and 1 in *SDHC)* [4]. Full details on the entire multi-institutional cohort can be found in **Supplementary Table 1.**

Despite superficially similar pathophysiology *(i.e.* the biallelic loss of a TCA metabolic enzyme), SDHRCC and FHRCC are molecularly distinct entities. While SDHRCC appears mainly to be driven by a germline mutation along with CNAs (loss of the wild-type chromosomal 1p arm), FHRCC demonstrates more genomic diversity. Of the 22 FHRCC that were consented to germline analysis and evaluated, 17 patients had a germline mutation, five of whom also had an additional somatic mutation (*i.e.* a composite mutation in *trans).* FHRCC patients without a germline mutation instead had a somatic mutation along with loss of chromosomal 1q arm, while none had purely somatic composite mutations [26]. Interestingly, a recent study published from our group noted that cases with biallelic somatic *FH* loss exhibited a similar clinical course to those with germline FH alterations [14]. We also observed a patient who presented with two somatic alterations to *SDHB* genes in the absence of germline mutation, resulting in the formation of a tumor with negative SDHB immunohistochemistry. This patient presented at an older age (55 years old) than the median of our cohort without a personal or family history of syndromic features associated with SDHRCC. To date, the patient has not developed metastasis after a follow up period of 53 months. To our knowledge, this is the first such report of purely somatic SDHB-deficient RCC [4,15,17,27]. This is novel as it expands our understanding of potential methods of inactivation for *SDH* genes. We further observed that SDHRCC tumors had lower TMB than FHRCC tumors as well as a decreased proportion of the fraction of the genome altered. Of the SDHRCC patients, 5 out of 11 (45%) had co-occurring mutations in cancer-associated genes (p = 0.05), notably *NF2* and *MTOR.* FHRCC tumors often had *FH* mutations co-occur with several other mutations such as *NF2, FAT1, PTPRT,* among others. 21 out of 25 (84%) FHRCC patients appeared to have co-occurring mutations in cancer-associated genes. The comparative genomic stability of SDHRCC relative to FHRCC suggests that, from an evolutionary perspective, SDHRCC tumors fail to genomically diversify and that biallelic loss of *SDHB* is sufficient to drive tumorigenesis.

Because complete loss of SDH or FH disrupts normally high metabolic flux through the TCA cycle, large scale changes to metabolism are expected. Although our cohort was limited due to the rarity of these tumors and the requirement for fresh-frozen tissue, metabolomic changes characterizing these diseases were evident. Grouping SDHRCC and FHRCC tumors together, there was a clear separation between SDHRCC/FHRCC compared to ccRCC as well as normal kidney. Although succinate was elevated in both SDHRCC/FHRCC compared to normal kidney tissue, fumarate was elevated exclusively in FHRCC. When SDHRCC/FHRCC tumors were compared against the normal kidney, we observed an extreme elevation of the purine derivative guanine. We suspect a mechanism unrelated to purine catabolism as there was no evident excess of xanthine, the catabolic breakdown product of guanine. This accumulation appeared pathognomonic, and was evident in a suspected SDHRCC found within our unclassified RCC cohort.

When comparing FHRCC and SDHRCC metabolomically, we observed that urea cycle metabolites such as argininosuccinate, citrulline, and fumarate were more elevated in FHRCC tumors when compared to normal kidney than in SDHRCC when compared to normal kidney [24]. Consistent with prior work [24], we hypothesize that this is due to a reversal of activity of argininosuccinate lyase in the urea cycle in the presence of an excess of fumarate. FHRCC tumors also had an elevation of N6-succinyladenosine compared to SDHRCC. A study performed in 2013 examined the urine of FH-deficient mice and found urine metabolites of urobilin, 2SC, fumarate and argininosuccinate consistently elevated [24], suggesting that these metabolites may be useful as non-invasive diagnostic biomarkers. Additionally, they investigated the use of an arginine-depleting enzyme, demonstrating that inhibiting the conversion of arginine to argininosuccinate reduced the proliferation of FH-deficient cells [24]. On the other hand, SDHRCC patients appeared to have an elevation of numerous acylcarnitine species compared to FHRCC. Mechanistically, SDH/Complex II is linked to beta oxidation of fatty acids through its role in FADH_2_ oxidation. This nominates fatty acid oxidation/FAD regeneration as both a biomarker and putative therapeutic target in SDHRCC.

On PCA analysis of metabolomic data, unclassified tumors co-localized with SDHRCC and FHRCC tumors. Of the four unclassified tumors, one tumor (RCC260) exhibited elevated levels of succinate and guanine without elevation of fumarate or N6-succinyladenosine. This metabolic profile resembled the 3 SDHRCC tumors that had been identified and diagnosed through genomic analysis or immunohistochemistry. Subsequently, IHC staining was performed on RCC260, which showed the loss of SDHB protein for this tumor. Thus, metabolomic profiling may help in the diagnosis, particularly in ambiguous cases, of SDHRCC. Prior genomic profiling of URCC tumors has shown several enriched molecular subsets within this heterogenous group, including those with *NF2* loss, mTOR activation, *FH* loss, and *ALK* translocation [28]. Given the overlap of some URCC cases with SDHRCC and FHRCC, future studies which phenotypically cluster URCC into metabolomic subsets may similarly uncover associations with clinical outcomes and novel targets for therapy.

The preferred systemic therapy for patients with SDHRCC and FHRCC is unknown, and treatments are largely extrapolated from experience with ccRCC as more prospective studies are performed. Uniquely, for FH-deficient patients, combination bevacizumab and erlotinib has shown significant clinical benefit in HLRCC patients (sporadic and syndromic) [29], and in a multi-institutional retrospective cohort, 3/5 FH-deficient patients had a partial response to cabozantinib monotherapy [29]. A retrospective study recently published from our group discussed response rates to systemic therapies in FHRCC patients [14]. Combination therapy targeting both mTOR and VEGF demonstrated the highest overall response rate (ORR = 44%) and disease control rate (DCR = 77%) amongst other systemic therapies such as VEGF monotherapy (ORR 20%, DCR 53%), checkpoint inhibitor monotherapy (ORR 0%, DCR 38%), and mTOR monotherapy (ORR 0%, DCR 25%) [14]. Although our ambition was to perform a similar kind of analysis for the SDHRCC cohort, a limited number of SDHRCC patients with available treatment information precluded such analysis (summarized in **Supplementary Table 2**). As SDHRCC becomes an increasingly recognized disease entity, clinical experience with common agents like VEGF, mTOR and immune checkpoint inhibitors will be better characterized in the future.

As highlighted in this study, metabolomic profiling provides additional data into tumor pathophysiology and may aid in phenotypic classification of tumors. As the field moves toward molecular stratification of RCCs [30], integrating metabolomic data may provide a new lens to distinguish tumor subtypes and uncover novel therapeutic targets. For example, the oncometabolite accumulation of fumarate and succinate through enzymatic loss of function has been shown to suppress homologous recombination thereby rendering tumor cells vulnerable to synthetic lethality with poly (ADP-Ribose) polymerase (PARP) inhibitors [31]. FH and SDH have also been shown to regulate HIF-alpha prolyl hydroxylases, thereby leading to stabilization and activation of HIF-1 alpha [32], [33]. Similar to the development and clinical activity seen with HIF2-alpha inhibitors [34], further study into the downstream effects of enzymatic activity loss may provide a mechanistic target for similar agents. Lastly, serum 2-hydroxyglutarate (2HG) has been isolated as a biomarker via peripheral blood [35], [36] and through radiographic detection [37] for isocitrate-dehydrogenase (IDH) mutant tumors. Given its similar role within the TCA cycle, future studies which explore fumarate, succinate, or other TCA associated metabolites may uncover novel dynamic biomarkers in this patient population. With the expanding utility of metabolomic analyses and viability of these studies on archival tissue compared to fresh-frozen tissue, metabolomic data may also be increasingly integrated into correlative studies of these rare tumor types.

Our study has limitations that are inherent when studying rare tumors. Adding to the challenge, the formal recognition of FHRCC and SDHRCC was only recently updated in 2016 [1]. To increase our sample size, we included patients from an outside multi-institutional cohort which had limited clinical information available. We also acknowledge that there is heterogeneity in the type and depth of genetic (targeted sequencing, WES, WGS) and metabolomic analyses. Despite the statistical limitations and heterogeneity of data and information, our genomic and metabolomic analysis of FHRCC and SDHRCC provides mechanistic insights into the defining molecular phenotypes of these entities and serve as a foundational resource for the broader research community.

In sum, while these tumor types fall under the umbrella of TCA-cycle mutant tumors, SDHRCC and FHRCC are genetically and metabolically distinct entities. Integration and application of genomic and metabolomic profiling in translational studies may enhance characterization of RCC tumor subtypes which share overlapping pathways in tumorigenesis, and may ultimately drive future discovery in biomarker and therapeutic discovery efforts.

## Methods

### MSK-IMPACT targeted sequencing

For patients on which targeted sequencing was done, tumor and matched-normal samples were sent for targeted sequencing using our previously-validated sequencing panel (MSK-IMPACT^®^), of which three versions exist, targeting 341, 410, or 468 actionable cancer-associated, respectively [18]. DNA from tumor and matched blood normal specimens were extracted from each patient and sheared to create barcoded DNA libraries. Using the captured DNA, all coding exons of the targeted genes, as well as a subset of polymorphic loci (for copy number analysis) were sequenced. Deep sequencing was performed at an average of 500X to 1,000X coverage. After alignment to the reference human genome, somatic alterations (including missense mutations, small insertions and deletions, structural rearrangements, and DNA copy number changes) were identified using a bioinformatics pipeline described in detail previously [18]. The oncology knowledge database OncoKB^®^ was used to annotate mutations to assess oncogenicity [38].

### Whole Exome Sequencing, processing, and mutation analysis

WES samples (from 8 SDHRCC patients; 2 with matched normals and 6 unmatched) were processed and analyzed using the TEMPO pipeline (v1.3, https://ccstempo.netlify.app/). In brief, demultiplexed FASTQ files were aligned to the b37 assembly of the human reference genome from the GATK bundle using BWA mem (v0.7.17). Aligned reads were converted and sorted into BAM files using samtools (v1.9) and marked for PCR duplicates using GATK MarkDuplicates (v3.8-1). Somatic mutations (single nucleotide variants and small insertions and deletions) were called in tumor-normal pairs using MuTect2 (v4.1.0.0) and Strelka2 (v2.9.10), and structural variants were detected using Delly (v0.8.2) and Manta (v1.5.0). Variants were annotated and filtered for recurrent artifacts and false positives using methods as previously described [39]. As with the IMPACT samples, OncoKB knowledge base was used to identify oncogenic alterations [38]. TMB was defined as the number of non-synonymous mutations in canonical exons per megabase. For copy number analysis, also as with the targeted sequencing data, we used FACETS version 0.5.6, described in detail below.

### Copy Number and Mutation Analysis

For zygosity determination, genome-wide total and allele-specific DNA copy number, purity, and ploidy were calculated via FACETS version 0.5.6 [40]. The expected number of copies for each mutation was generated based on observed variant allele fraction and local ploidy [41]. Cancer cell fractions were calculated using a binomial distribution and maximum likelihood estimation normalized to produce posterior probabilities [42]. We then assessed what fraction of samples have either gains or LOH (either absolute loss of copy or cnLOH) at each cytoband in the genome, applying a clonality cutoff of 0.25. In generating the oncoprints, we restricted the gene set to genes that are mutated at least 3 times in our sample set, and classified mutation events (instances where a gene is altered in a patient) according to the type of mutation and whether there were multiple mutations. We compared tumor mutation burden (TMB) of FHRCC with that of SDHRCC by taking the total number of non-silent somatic mutations across IMPACT-341 genes for patients in each subtype and applying the Wilcoxon test to assess whether the distributions were significantly different. To compare fraction genome altered (FGA), we computed the fraction of the sequenced portion of the genome (which varies depending on whether the sample is WES, WGS, or IMPACT) subject to either loss or gain of copy or copy-neutral LOH. We then compared FGA between FHRCC and SDHRCC using the Wilcoxon test.

### Metabolomic Profiling

After acquiring informed consent and Memorial Sloan Kettering Cancer Center institutional review board approval, partial or radical nephrectomies were performed at Memorial Sloan Kettering Cancer Center (New York) and stored at the MSK Translational Kidney Research Program (TKCRP). Samples were flash frozen and stored at −80 degrees Celsius prior to metabolomic characterization. Clinical metadata was recorded for all tumor samples. Samples were thawed and extracted according to Metabolon’s standard protocol, which removed proteins, dislodged small molecules bound to the protein or physically trapped in the protein matrix, and recovered a wide range of chemically diverse metabolites. Samples were then frozen, dried under vacuum and prepared for LC/MS.

The sample extract was split in two and reconstituted in acidic and basic LC-compatible solvents. The acidic extracts were gradient eluted using water and methanol containing 0.1% Formic acid, while the basic extracts, which also used water and methanol, contained 6.5 mM Ammonium Bicarbonate. One aliquot was analyzed using acidic positive ion optimized conditions and the other used basic negative ion optimized conditions. The aliquots were two independent injections using separate dedicated columns. The MS analysis alternated between MS and data-dependent MS/MS scans using dynamic exclusion. The LC/MS portion of the platform was based on a Waters ACQUITY UPLC and a Thermo-Finnigan LTQ-FT mass spectrometer, which had a linear ion-trap (LIT) front end and a Fourier transform ion cyclotron resonance (FT-ICR) mass spectrometer backend. Accurate mass measurements could be performed for ions with counts greater than 2 million. The average mass error was less than 5 ppm. Ions with less than 2 million counts required a greater effort to characterize. Typically, fragmentation spectra (MS/MS) were generated in a data-dependent manner, but targeted MS/MS could be employed if necessary, such as in the case of lower level signals.

Data was extracted from the raw mass spectrometry files, which was loaded into a relational database. The information was then examined and appropriate QC limits were imposed. Metabolon’s proprietary peak integration software was used to identify peaks, and component parts were stored in a separate data structure. Metabolites were compared to an in-house library of standards from Metabolon. Data on each of these standards were based on retention index, mass-to-charge ratio, and MS/MS spectra. These parameters of each feature for each compound in the metabolomic data were compared to analogous parameters in the library. As described in Evans et al. [43], compounds were identified based on three criteria: retention index within 75 RI units of the proposed identification, mass within 0.4 m/s and MS/MS forward and reverse match scores. Each compound was corrected in run-day blocks by registering the medians to equal one and normalizing each data point accordingly. The data was subsequently log_2_ normalized. When metabolite levels were below the level of detection, the lowest measured abundance of that compound across all samples was imputed. All statistical tests were performed in R. Tests comparing distributions were performed using wilcox.test. All statistical analyses were two-sided and p-values were Benjamini-Hochberg corrected.

## Acknowledgements

We thank the members of the Reznik and Hakimi laboratories for discussion and support. We additionally thank Divya Bezwada for keen insights and critical feedback.

---

**Supplementary Figure 1:**
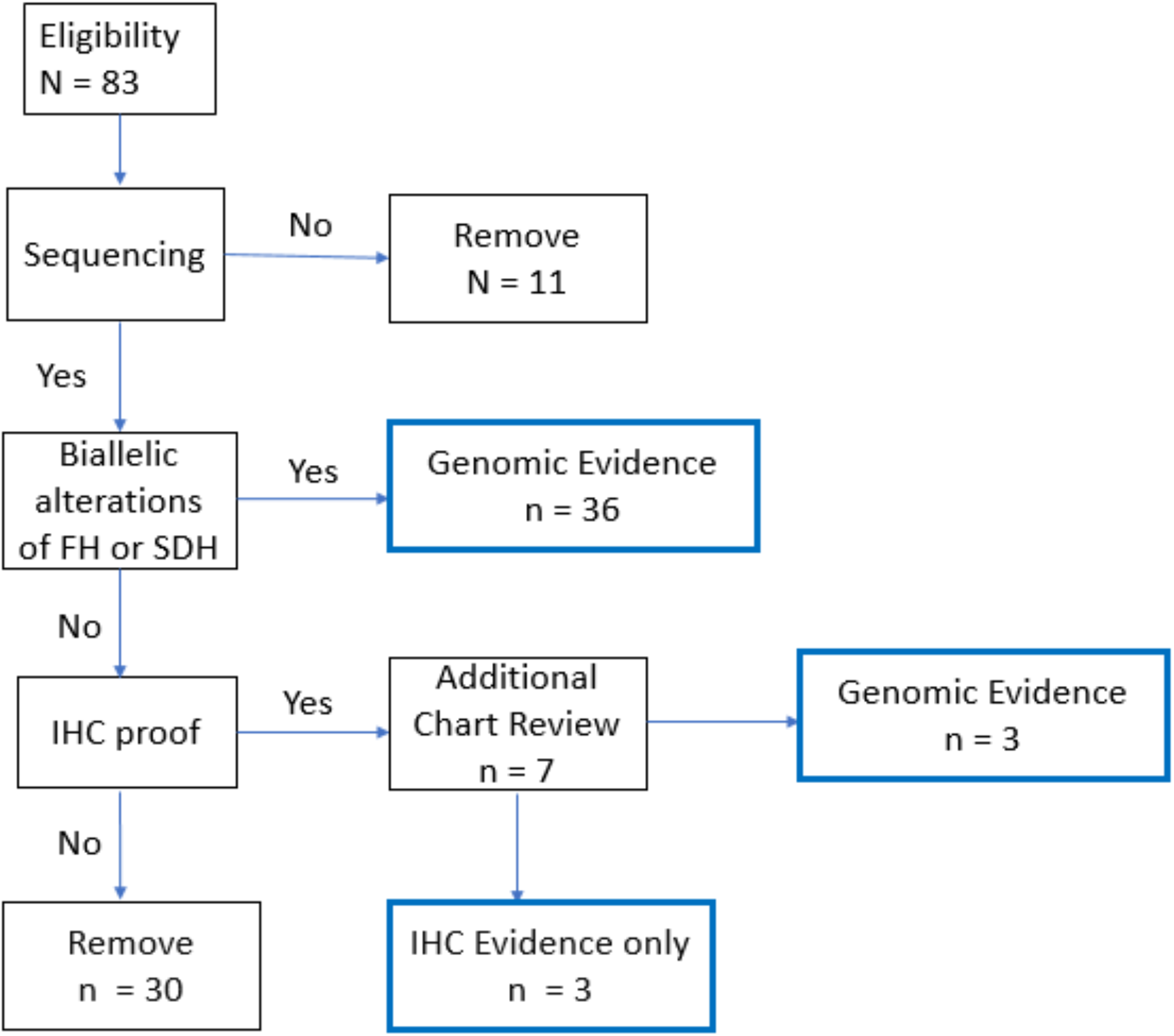
83 RCC patients with a presumed diagnosis of FHRCC or SDHRCC were identified using a combination of genomic, immunohistochemical (IHC), and expert genitourinary pathology review. Of these, 11 patients were excluded due to lack of sequencing data either through MSK-IMPACT, WES or WGS. Patients who demonstrated biallelic alterations of *SDHA/SDHB/SDHC/SDHD/SDHAF2* or *FH* were considered to have genomic evidence of disease. 36 patients demonstrated biallelic loss of the gene of interest. For the remaining patients, immunohistochemical staining was evaluated for loss of protein expression (FH or SDH). 6 patients demonstrated loss of FH or SDH on IHC, prompting additional chart review for genomic evidence of disease. Of the 6 that were evaluated, 3 patients were identified as having biallelic alterations due to genetic testing either at an outside institution or through a different clinical genetic service. 3 patients IHC evidence of disease only. The final cohort consisted of a total of 42 patients, 25 patients were FH-deficient and 17 patients were SDH-deficient.

## Notes

**Disclosure of Conflicts of Interest**, Chung-Han Lee reports personal fees from Amgen, BMS, Exelixis, Eisai, Merck, Pfizer, and EMD Serono outside the submitted work. Robert J. Motzer reports serving in a consultancy or advisory role for Pfizer, Novartis, Genentech/Roche, Lilly Oncology, Eisai and Exelixis, and received research funding from Bristol-Myers Squibb, Pfizer, Eisai and Exelixis. Victor E. Reuter is a non-compensated advisor for PAIGE.AI. A. Ari Hakimi reports Advisory Board compensation from Merck. No disclosures were reported by the other authors.

### Competing Interest Statement

Chung-Han Lee reports personal fees from Amgen, BMS, Exelixis, Eisai, Merck, Pfizer, and EMD Serono outside the submitted work.
Robert J. Motzer reports serving in a consultancy or advisory role for Pfizer, Novartis, Genentech/Roche, Lilly Oncology, Eisai and Exelixis, and received research funding from Bristol-Myers Squibb, Pfizer, Eisai and Exelixis.
Victor E. Reuter is a non-compensated advisor for PAIGE.AI.
A. Ari Hakimi reports Advisory Board compensation from Merck.
No disclosures were reported by the other authors.

